# DOC Release and Physiological Response of *Zostera marina* Under Light and Nutrient Gradients

**DOI:** 10.64898/2026.05.22.726822

**Authors:** Lizhi Cheng, Tian Li, Xiaojie Zhao, Yuzhang Zhu, Wentao Wang, Yiwei Wu, Caiyan Xv, Mingliang Zhang, Bin Zhou, Rongyu Xin, Huawei Qin

**Affiliations:** College of Marine Life Science, Ocean University of China, Qingdao, 266003, China; Marine Carbon Sink Research Center, Shandong Marine Resource and Environment Research Institute, Yantai 264006, China; College of Fisheries and Life Science, Shanghai Ocean University, Shanghai, 201306, China

**Keywords:** *Zostera marina*, Seagrass Bed, Dissolved Organic Carbon (DOC), light, Eutrophication

## Abstract

The seagrass bed, known as an important blue carbon ecosystem, not only has robust carbon sequestration capacity, but also releases the fixed organic carbon into the water column in the form of dissolved organic carbon (DOC). *Zostera marina* is the main constructive species of the seagrass meadows of northern China, but the regulatory mechanism of its DOC release in response to environmental changes still remains unclear. Adult *Z. marina* was used as the experimental material in this research to investigate the effects of light intensity and nutrient concentration on its DOC release, and to observe its physiological responses. By using a total organic carbon (TOC) analyzer, dissolved oxygen monitoring, transmission electron microscopy (TEM), and transcriptome analysis, we explored how light and nutrients affect DOC release and the underlying physiological mechanisms in *Z. marina*. The results showed that both elevated light intensity and short-term nutrient enrichment promoted organic carbon release from *Z. marina* to the environment. Increased light enhanced the net primary productivity (NPP) of *Z. marina*, while the magnitude of DOC release increased more than NPP itself. Nutrient enrichment reduced NPP while raising the proportion of DOC released by *Z. marina*. Transcriptome analysis showed that nutrient enrichment significantly affected pathways, including flavonoid biosynthesis and fatty acid degradation. This study demonstrated that changes in light and nutrients can significantly influence DOC release and carbon allocation patterns in *Z. marina*, providing a theoretical basis for blue carbon accounting and ecological restoration evaluation in seagrass beds.

## 1. INTRODUCTION

Blue carbon refers to the carbon that is captured by the oceans, accounting for more than 55% of green carbon. Carbon caught by marine organisms can be stored in the sediments of mangrove communities, salt marshes, and seagrass beds over long timescales measured in millennia(Nellemann et al., 2009). Carbon fixed by blue carbon ecosystems mainly exists as particulate organic matter (POM) and dissolved organic matter (DOM) in the water column and participates in carbon sedimentation via the microbial carbon pump(MCP) (Jiao et al., 2010). DOM is defined as organic matter that passes through a 0.45 μm filter (or equivalent), and its carbon content is called dissolved organic carbon (DOC), while the carbon retained on the filter is particulate organic carbon (POC) (Potter and Wimsatt, 2009). DOC represents the second-largest carbon pool and the largest organic carbon pool in the ocean, with a standing stock of approximately 662 Pg carbon (Hansell and Carlson, 2013), far exceeding that of POC, making it a key focus in blue carbon research. Based on biodegradability, it can be broadly divided into labile DOC (LDOC) and recalcitrant DOC (RDOC). The high abundance of RDOC is critical to making the marine DOC pool the second-largest global carbon reservoir (Hansell et al., 2012) and is closely linked to blue carbon sequestration. (Jiménez-Ramos et al., 2022a; Zhao et al., 2024) Therefore, exploring the release of LDOC production and RDOC transformation is an important direction in blue carbon research. Current studies on oceanic organic carbon production have mostly focused on marine algae and coral communities. (Naumann et al., 2010; Hertkorn et al., 2013; K. Wang et al., 2024; N. Xu et al., 2022), while research on seagrasses, especially their DOC release mechanisms, remains relatively scarce and mechanistically incomplete(Egea et al., 2019).

As one of the three major blue carbon ecosystems in the ocean, seagrass meadows play a vital role in global carbon storage(Fourqurean et al., 2012; Serrano et al., 2021; Fu et al., 2023). Although they cover less than 0.1% of the world’s ocean surface, they account for 13.3% of global coastal net ecosystem productivity and store 12.7% of organic carbon(Duarte et al., 2005). However, the carbon sequestration capacity of seagrass beds may still be underestimated(Duarte and Krause-Jensen, 2017). Findings from studies on lateral carbon fluxes in seagrass, mangrove, and coral communities and their adjacent areas further support this view: seagrass ecosystems act as net sources of all carbon forms, creating an important pathway for carbon release from sediments or plant tissues into the water column. But carbon inputs from mangroves and rivers delivered laterally into seagrass meadows can partially offset their potential carbon sink capacity(Akhand et al., 2021).Therefore outdoor studies of seagrass ecosystems may underestimate the c carbon sink function of seagrass. Overall, seagrass ecosystems are crucial in blue carbon fixation. Seagrasses form extensive meadows in shallow (Larkum et al., 2006), sheltered coastal waters, providing key ecosystem services including sediment stabilization, habitats and nurseries, nutrient cycling, and material transport(Fourqurean et al., 2012), and represent an important component of blue carbon ecosystems and a priority for restoration(Sanders et al., 2026).

Within seagrass bed ecosystems, DOC is the most dynamic carbon pool (Krause et al., 2025), with multiple sources including release during sediment resuspension (Dahl et al., 2020), release by seagrasses themselves(Kaldy, 2012), transformation via the MCP(Dahl et al., 2020), and advective exchange with external water masses(Egea et al., 2023). This paper primarily investigates the DOC released by seagrasses themselves. Two major hypotheses for DOC release by aquatic primary producers have been proposed based on key environmental drivers: the light-dependent overflow mechanism(Fogg, 1983) and the nutrient-driven diffusion mechanism(Bjørnsen, 1988). The former held that enhanced photochemical reactions under high photon flux could increase DOC release(Fogg, 1983; Egea et al., 2019). The latter held that the promoted synthesis of chlorophyll(Carstensen et al., 2018), NADPH, ATP, and the accelerated regeneration of ribulose-1,5-bisphosphate (RuBP)(J. Li et al., 2022), could stimulate the photosynthetic cycle and thus improve carbon fixation efficiency and DOC release(Bjørnsen, 1988; Liang et al., 2020; Bao et al., 2022).

The growth of seagrass beds is closely linked to ambient light intensity: the light requirement of seagrasses is usually characterized by the percentage of surface incident light, which increases with the rise of water turbidity and sediment organic matter content, which directly influences the selection of sites and timing for seagrass transplantation. (Kenworthy et al., 2014; S. Xu et al., 2020). Controlled experiments have also demonstrated that seagrasses exhibit clear minimum light thresholds and light stress responses(Bertelli and Unsworth, 2018). Meanwhile, nutrient availability in the marine environment is closely related to seagrass growth. Previous studies have demonstrated that high-nutrient conditions can accelerate the mineralization of RDOC in seagrass ecosystems(Liu et al., 2020), which has also been further confirmed by laboratory incubation experiments(Zhang et al., 2025). However, the effects of high nutrients on seagrass growth itself and DOC release still remain unclear. Under the policy background of Dual Carbon, clarifying the environmental regulation of seagrass DOC release is of practical significance for blue carbon accounting and environmental restoration assessment. there remains a lack of systematic evidence regarding the regulation of DOC efflux in *Z. marina* when both factors are present simultaneously(J. Kaldy, 2012; J. E. Kaldy, 2014), there remains a lack of systematic evidence regarding the regulation of DOC efflux in *Z. marina* when both factors are present simultaneously. Based on the above research background and gaps, our study selected Z. marina, a constructive species in temperate coastal areas, as the research object. Under controlled laboratory conditions, two single-factor gradient experiments of light intensity and nitrogen and phosphorus nutrients (NH^4+^, PO_4_^3-^), a two-factor orthogonal experiment, and a short-term day-night alternating culture experiment were conducted. We hypothesized that there were significant differences in the dominant controlling factors and mechanisms under the DOC release process of *Z. marina* under different light intensities and nutrient concentrations. This study will verify and refine the relevant mechanisms at the microscale, aiming to deepen the understanding of the DOC release process of *Z. marina*, and provide a theoretical basis and scientific reference for blue carbon accounting and ecological restoration of seagrass meadows.

## 2. MATERIAL AND METHOD

### 2.1 Collection and temporary cultivation of research subjects

The *Z. marina* used in the experiment was collected from the intertidal zone of Zhanqiao Pier, Qingdao, China (120°19′20″E, 36°3′39″N, WGS-84) in July-August 2025.

Salinity of the ambient seawater was measured as 33‰ using a multiparameter probe (YSI Pro Quatro, Xylem Inc., USA) during *Z. marina* collection. Meanwhile, we used 10 BKMAMLAB 50 mL centrifugal tubes to collect in-situ seawater for ammonia nitrogen and total phosphorus determination(Ahmed et al., 2023). The results showed that the average content of ammonia nitrogen was 0.20 mg·L^−1^, and the average content of total phosphorus was 0.0233 mg·L^−1^. The artificial seawater used in subsequent experiments had a salinity of 33‰, an average ammonia nitrogen concentration of 0.21 mg·L⁻¹, and an average total phosphorus concentration of 0.0257 mg·L⁻¹

The Mann-Whitney U test was used to conduct a nonparametric statistical analysis on the salinity, ammonia nitrogen, and total phosphorus of artificial seawater and the water sample from the collection site. The result showed that there was no significant difference in ammonia nitrogen (p = 0.591) and phosphorus (p = 0.354) between the two groups, indicating homogeneity in statistical terms. Hence, it can be concluded that the in-situ seawater collected from the Zhanqiao Pier has good consistency with artificial seawater in basic properties, and the artificial seawater can be used as an alternative for ambient seawater in the experiment.

Epiphytic algae and mollusks were gently removed from the collected *Z. marina* using soft brushes prior to use(Monteiro Vasconcelos et al., 2024). The rhizomes were then trimmed to a length of 2 cm for each individual. The cleaned *Z. marina* was placed in a 120 L polyethylene tank and cultivated using artificial seawater until the experiments.

### 2.2 Experimental design

#### 2.2.1 DOC release experiment under nutrient gradients

The *Z. marina* was cultured in illumination incubators (Zhujiang Brand, LRH-800-G, China) and illumination incubators (Ningbo Jiangnan Instrument Factory, GXZ-500D, China), with light intensity adjusted by blackout fabrics. Acid-washed glass containers (17 cm × 17 cm × 30 cm, wall thickness = 0.7 cm) were used for cultivation. The *Z. marina* was incubated at 18 °C with a 12h light cultivation, and the light intensity set was 100 (94.97 ± 6.47) μmol· m^−2^ · s^−1^(Dennison and Alberte, 1985). Nutrient gradients were defined by dissolved inorganic nitrogen (DIN) with NH_4_ ^+^ as the nitrogen source and by dissolved inorganic phosphorus (DIP) with PO_4_ ^3-^as the phosphorus source. Ammonia nitrogen concentration was supplemented using BKMAMLAB NH_4_Cl (batch No. 20241225, 2025.3.15), while phosphate concentration was supplemented using SCR Na_3_PO_4_·12H_2_O (batch No. 20240920, 2025.3.29). The experimental DIN and DIP gradients were established with reference to Chinese national and coastal eutrophication assessment standards, including GB 3097—1997 and T/CSES 174—2024, and were further informed by international eutrophication assessment frameworks such as OSPAR and HELCOM (State Environmental Protection Administration of China and State Bureau of Quality and Technical Supervision, 1997; OSPAR Commission, 2022; Chinese Society for Environmental Sciences, 2024; HELCOM, 2026). The DIN ranged from 2.4∼9.6 mg·L^−1^ (≈187-621 μM), while the DIP ranged from 0.24∼0.96 mg·L^−1^ (≈ 9-31 μM). Ammonia nitrogen and total phosphorus were measured using a multiparameter water quality analyzer (LH-M300, Lianhua Tech, China). Light intensity was monitored using a photosynthetically active radiation meter (HPL-220P, HopooColor, China). The results were shown in supplementary table1 (ST1). The wet weight of each replicate was maintained between 70 and 80 g.

#### 2.2.2 DOC release experiment under light gradients

We built up four gradients: 50 μmol·m^−2^·s^−1^ (low light), 100 μmol·m^−2^·s^−1^ (standard light), 200 μmol·m^−2^·s^−1^ (high light), 350 μmol·m^−2^·s^−1^ (ultra high light) (Hasler-Sheetal et al., 2016). The culture conditions and parameters were the same as 2.2.1. The results of the measured light intensity were shown in supplementary table 2 (ST2). As the water for cultivating is artificial water collected from the same PE tank, we did not analyze the nutrient concentration.

#### 2.2.3 The two-factor orthogonal experiment

An ultra-high light intensity (350 μmol m^−2^ s^−1^) was applied, with ammonia nitrogen and total phosphorus concentrations set at 4.8 and 0.48 mg/L, respectively. The nutrient supplement and the method for measuring the cultivation conditions were the same as those in 2.2.1. Four groups were set up: wild type (WT), high-light treatment (HL), high-nutrients treatment (HN), and high-light and high-nutrients treatment (HLHN). The culture conditions and parameters were the same as 2.2.1. Results of the measured light intensity and nutrients were shown in supplementary table 3 (ST3).

#### 2.2.4 Short-term diurnal alternating culture experiment

We set up three groups: wild type (WT), high-light treatment (HL), and high-nutrient treatment (HN). The light intensity for HL, the ammonia nitrogen, and total phosphorus for HN were set at 300 μmol·m^−2^·s^−1^, 4.8 mg/L, and 0.48 mg/L, respectively. Results of the measured light intensity and nutrients were shown in supplementary table 4 (ST4). To ensure the *Z. marina* activity in the dark experimental phase as much as possible, we used the NETLEA aeration pumps for oxygen supply, and to maintain similar aeration rates among the 15 containers(Liu et al., 2023). Put 40-50 g of *Z. marina* into each container, then incubated the *Z. marina* at 18 °C with a 12h: 12h light: dark cycle. Blot the *Z. marina* surface and measured the wet weight before and after cultivation using a ZG-TP203 electronic balance. The wet weight of the *Z. marina* before the experiment was shown in supplementary table 4 (ST4).

### 2.3 Sample collection

#### 2.3.1 Collection of the culture water sample

We used the BKMAMLAB 0.45 μm glass fiber membrane, Syringe Filter, to collect DOC from the culture water sample(Herbert et al., 1993). Before collecting, the membranes were rinsed with ultra-pure water for three times(Lian et al., 2021; Tisserand et al., 2024). After collecting, we used an Eppendorf 5 mL pipette to aliquot the samples into three 20 mL amber sample vials for TOC measurement, and three 50 mL amber vials were collected for nutrient analysis. The pH of TOC measurement samples was adjusted to <2 with hydrochloric acid. All samples were stored at −20 °C until analysis.

The amber sample vials were rinsed with ultra-pure water twice, covered with Labshark aluminum foil, and then put into the FAITHFUL ceramic fiber muffle furnace together for a pre-combustion at 450 °C for 4 hours in order to remove residual organic carbon from the vials(Fourquez et al., 2025). A portable dissolved oxygen meter (Lei-ci, JPB-607A, China) was used to determine the concentration of dissolved oxygen.

#### 2.3.2 Collection of the TEM samples

The samples we used were from the short-term day-night alternating culture experiment mentioned in 2.2.4. After measuring the *Z. marina*’s wet weight, we randomly select leaves from each culture tank and cut four segments of 0.5 cm × 0.25 cm from each group. The segments were rinsed successively with ultrapure water and 1× PBS buffer, immediately transferred to 4% glutaraldehyde fixative, and subjected to vacuum infiltration to ensure complete tissue penetration. Samples were then fixed at 4 °C until examination.

#### 2.3.3 Collection of transcriptome sample

Leaf sampling and cleaning procedures were consistent with those mentioned in Section 2.3.2. Segments of 0.7 cm × 0.7 cm were excised from the same relative position of the leaves, with eleven samples per group. The samples were snap-frozen in liquid nitrogen immediately after collection and stored at −80 °C for subsequent transcriptome sequencing.

### 2.4 Sample detection

#### 2.4.1 TOC determination

DOC concentrations in water samples were measured using a TOC-L total organic carbon analyzer (Shimadzu, Japan) equipped with an ASI-L autosampler, following the non-purgeable organic carbon (NPOC) method(Stegen et al., 2025), at the Large Instrument Platform, College of Environmental Science & Engineering, Laoshan Campus, Ocean University of China, Qingdao, China. Calibration standards were prepared with ultrapure water. Each sample was measured in triplicate.

#### 2.4.2 TEM imaging

The fixed samples from Section 2.3.2 were rinsed three times with 0.1 M phosphate buffer (15 min each), post-fixed by 1% osmic acid for 1-1.5 h, and rinsed three times again. The samples were then dehydrated in a graded ethanol series(50%, 70%, 90% for 15 min each; 100% for 15-20 min, repeated three times), followed by gradual infiltration with Epon 812 resin at 37 °C, 45 °C, and 60 °C. The embedded samples were sectioned into approximately 70 nm ultra-thin slices using an ultramicrotome (Richert-Jung, ULTRACUT E, Austria). After staining with uranium acetate and lead citrate, the sections were observed and photographed under a transmission electron microscope (JEOL, JEM-1400F, Japan).

#### 2.4.3 Transcriptome extraction and sequencing

Total RNA was extracted from tissue samples using the TRIzol method. RNA concentration, purity, and integrity were evaluated using NanoDrop, agarose gel electrophoresis, and an Agilent Bioanalyzer. The mRNA was enriched with Oligo(dT) magnetic beads, followed by fragmentation, cDNA synthesis, adapter ligation, and PCR amplification, to construct the cDNA library. After library quantification, sequencing was performed on the Illumina NovaSeq platform. Fastp was used for quality control of sequencing data. HISAT2 was used for alignment to the reference genome. RSEM was used for gene expression quantification. DESeq2 was used for differential expression analysis. KEGG annotation and enrichment analysis were subsequently conducted.

### 2.5 Data analysis

All data were processed and statistically analyzed in GraphPad Prism 9.5. For each experiment, n represented the number of independent samples; the specific values were shown in the respective figures. Due to the small sample size in some experiments, the normality test was not fully effective, so the selection of statistical methods was mainly based on experimental design, data structure, and sample size.

We chose a two-tailed t-test for intergroup comparisons. For repeated measurements of the same sample before/after treatment, we chose a paired two-tailed t-test. For multiple comparisons with one independent variable, one-way ANOVA was used. The two-way ANOVA was used to analyze the two-factor orthogonal experiment.

The restricted maximum likelihood (REML) in Prism was used to handle repeated measurements and missing values in order to analyze repeated measures structure and allow the missing values to exist. When the spherical assumption was violated, the Geisser-Greenhouse correction was used to correct the degrees of freedom. Appropriate robust methods were adopted based on the homogeneity of variance if possible, with detailed explanations provided in the results. Post hoc multiple comparisons were performed using Tukey, Sidak, or Dunnett tests when necessary.

All tests were two-tailed, and *P* < 0.05 was considered statistically significant. Unless otherwise stated, data are presented as Mean ± SD.

## 3. RESULTS

### 3.1 Effects of light and nutrient salt gradients on the allocation of net primary productivity and oxygen production

Under different light intensity treatments (PPFD 50, 100, 200, 350), the DOC/NPP ratio (Fig.1a) showed significant differences among groups (one way ANOVA: *F*(3, 8) = 14.16, *P* = 0.0015), and the homogeneity of variances was satisfied (Brown-Forsythe test: *F*(3, 8) = 0.8921, *P* = 0.4858). Tukey’s post hoc test revealed that the DOC/NPP ratio in the PPFD = 350 treatment (20.11%) was significantly higher than that in the PPFD = 50 (11.80%) and PPFD = 200 (11.38%) treatments (both *P* < 0.01), while the PPFD = 100 treatment (15.33%) showed no significant differences with other treatments (*P* > 0.05), which was consistent with the letter grouping in the figure (350: a; 50 and 200: b; 100: ab).

**Fig. 1.**
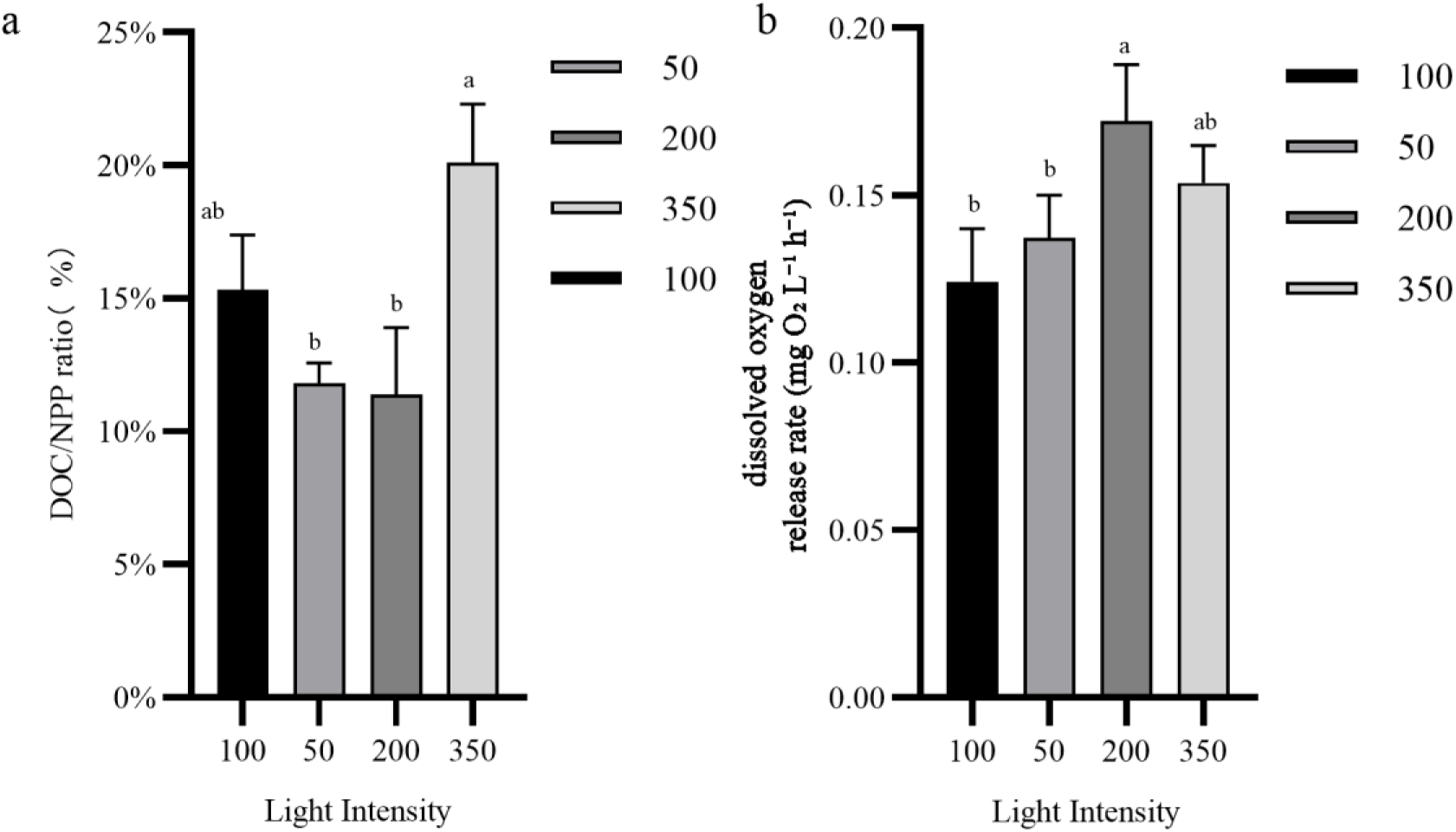
Carbon allocation and oxygen production responses under different PPFD levels (mean ± SD) (A) Ratio of dissolved organic carbon to net primary productivity (DOC/NPP ratio, %). (B) Hourly dissolved oxygen release rate (mg O2·L^−1^·h^−1^) Bars represent mean values, and error bars indicate standard deviations (mean ± SD). Different lowercase letters indicate significant differences according to Tukey’s post hoc multiple comparison test (*P* < 0.05), whereas the same letter indicates no significant difference. Light intensity is expressed as PPFD (50, 100, 200, and 350).

The hourly dissolved oxygen release rate (Fig.1b) also varied significantly with light intensity (one-way ANOVA: *F*(3, 12) = 8.403, *P*=0.0028), and the homogeneity of variances was satisfied (Brown-Forsythe test: *F*(3, 12) = 0.9762, *P* = 0.4362; Bartlett test: *P* = 0.9066). Tukey’s post-hoc test indicated that the dissolved oxygen release rate was the highest in the PPFD = 200 treatment (0.1723 mg O_2_·L^−1^·h^−1^), which was significantly higher than that in the PPFD = 100 (0.1242 mg O_2_·L^−1^·h^−1^) and PPFD = 50 (0.1373 mg O_2_·L^−1^·h^−1^) treatments (*P* < 0.01 and *P* < 0.05, respectively); the PPFD = 350 treatment (0.1538 mg O_2_·L^−1^·h^−1^) showed no significant differences with other treatments (*P* > 0.05) and presented an intermediate level. This result was consistent with the letter labels in the figure (200: a; 100 and 50: b; 350: ab).

The DOC/NPP ratio (Fig.2a) showed extremely significant differences among groups (one-way ANOVA: *F*(3, 12) = 79.80, *P* < 0.0001), and the homogeneity of variances was satisfied (Brown-Forsythe test: P = 0.2907). Tukey’s post hoc test revealed that the WT treatment had the lowest mean DOC/NPP ratio (10.38%), which was significantly lower than the 2.4 (37.99%),4.8 (51.80%), and 9.6 (52.41%) treatments (all *P* < 0.0001). Meanwhile, the 2.4 treatment was also significantly lower than the 4.8 and 9.6 treatments (*P* = 0.0039 and 0.0028, respectively, after adjustment), while no significant difference was observed between the 4.8 and 9.6 treatments (*P* = 0.9972). Overall, the DOC/NPP ratio increased significantly with increasing nutrient concentration and reached a plateau at 4.8-9.6, which was consistent with the letter grouping in the figure (WT: c, 2.4: b, 4.8 and 9.6: a).

**Fig. 2.**
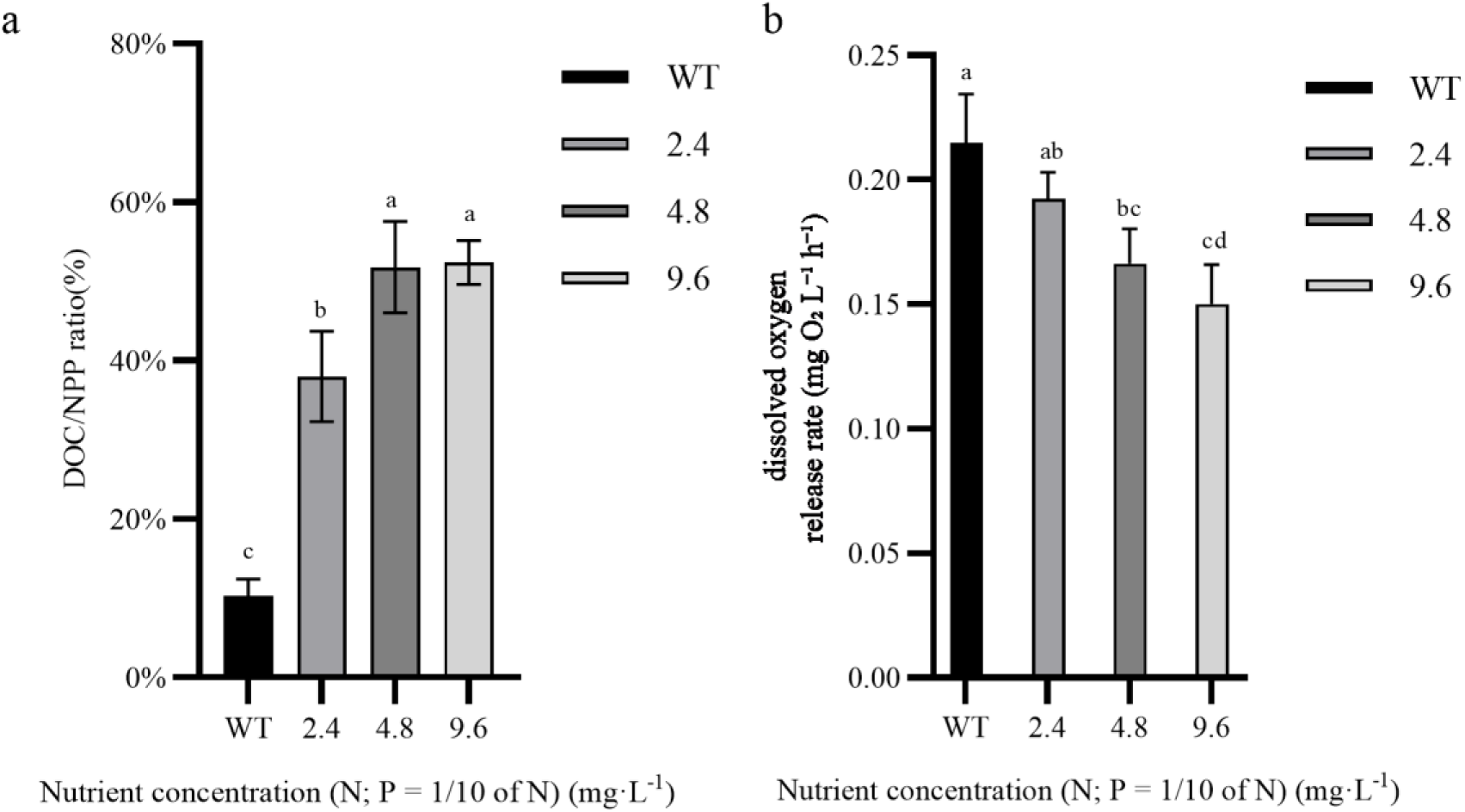
Carbon allocation and oxygen production responses under single-gradient nutrient concentrations (N, P = 1/10 N)(mean ± SD) (A) Ratio of dissolved organic carbon to net primary productivity (DOC/NPP ratio, %). (B) Hourly dissolved oxygen release rate (mg O_2_·L^−1^·h^−1^) Bars represent mean values, the error bars indicate standard deviation (mean ± SD). Different lowercase letters indicate significant differences in Tukey’s post hoc multiple comparisons (*P* < 0.05), while the same letter indicates no significant difference. The x-axis shows nutrient concentration gradients (WT, 2.4, 4.8, 9.6; unit mg·L-1, expressed as N, and phosphorus (P) concentration is 1/10 of N).

The hourly dissolved oxygen release rate (Fig.2b) also showed significant differences among groups (one-way ANOVA: *F*(3,12) = 13.79, *P* = 0.0003). The results of homogeneity of variance tests indicated that the data satisfied the homogeneity of variance assumption (Brown-Forsythe test: *F*(3, 12) = 0.4613, *P* = 0.7145; Bartlett test: *P* = 0.8082). Tukey’s post hoc test revealed that there were significant differences existing between the WT treatment and the 4.8 (*P* = 0.0037) and 9.6 (*P* = 0.0003) treatments; a significant difference was also observed between the 2.4 and 9.6 treatments (*P* = 0.0101). In contrast, no significant differences were found between all adjacent treatment groups (all *P* > 0.05). Overall, as the nutrient concentration increased, the hourly dissolved oxygen release rate showed a decreasing trend, with the 9.6 treatment having the lowest rate and the WT treatment having the highest rate, indicating that higher nutrient levels may exert an inhibitory effect on dissolved oxygen release, which was consistent with the grouping trend of shared letter labels among partial treatments in the figure.

### 3.2 Interactive effects of light and nutrient gradients on the allocation of net primary productivity and oxygen production

In the two-factor orthogonal experiment of light and nutrient gradients, the DOC/NPP ratio (Fig.3a) was significantly affected by the main effects of nutrients and light. Two-way ANOVA analysis showed that the main effect of nutrients was extremely significant (*F*(1, 12) = 54.11, *P*<0.0001), explaining 73.02% of the total variation; the main effect of light was also significant (*F*(1, 12) = 5.435, *P* = 0.0380), explaining 7.33% of the variation; meanwhile the light×nutrient interaction was not significant (*F*(1, 12) = 2.557, *P* = 0.1358). A comparison of the mean values showed that the mean DOC/NPP ratio under high light conditions was higher than that under standard light (74.90% vs 58.26%), and high nutrients significantly increased the DOC/NPP ratio (92.83% vs 40.33%). After Sidak post-hoc correction, high nutrients were significantly higher than standard nutrients under both standard light and high light (e.g., Std nutrients-Std light vs High nutrients-Std light: *P* = 0.0093; Std nutrients-High light vs High nutrients-High light: *P* = 0.0002), while no significant difference was observed between standard light and high light at the same nutrient level (Std nutrients: *P* = 0.9967; High nutrients: *P* = 0.0960).

**Fig. 3.**
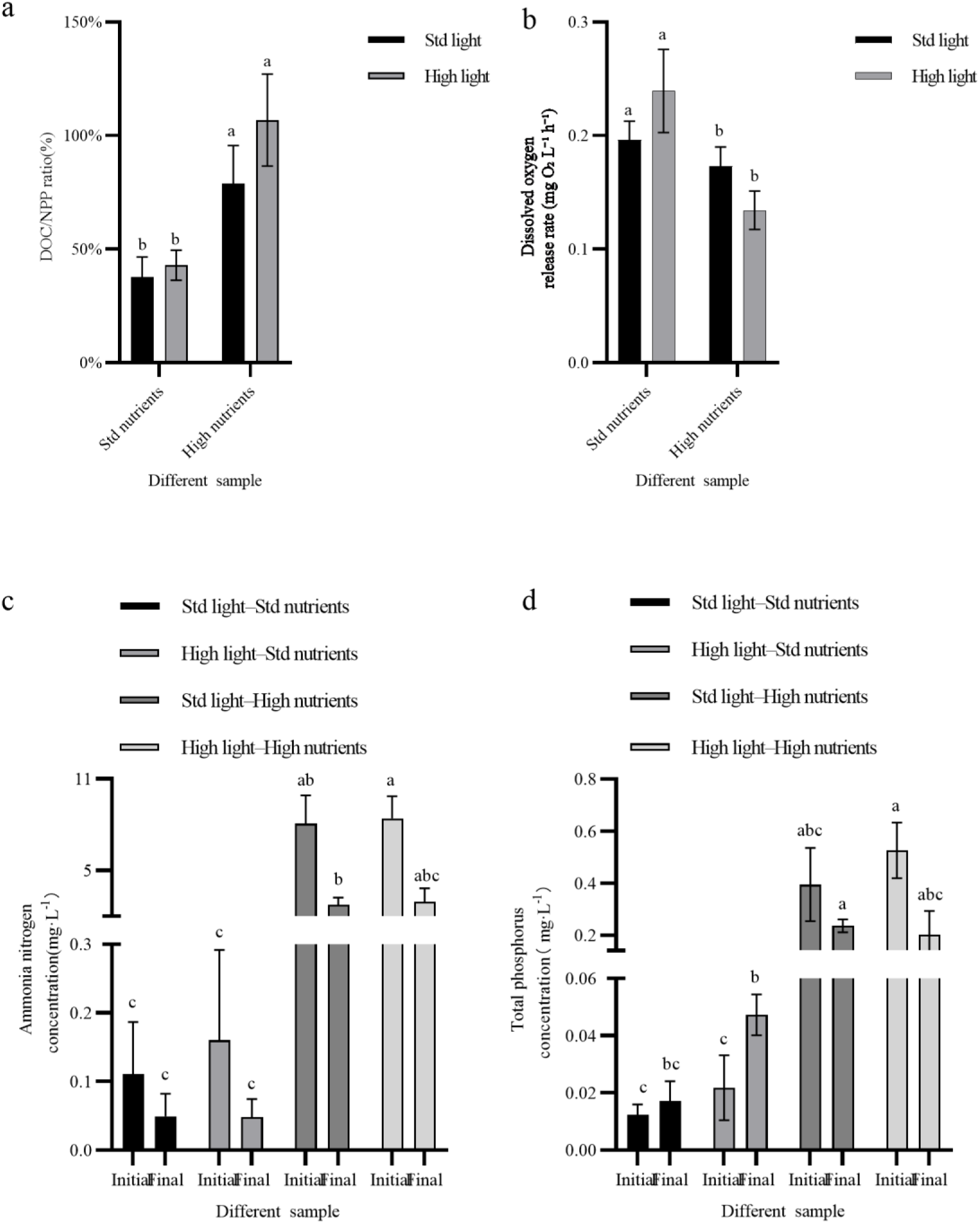
Carbon allocation and oxygen production responses under single-gradient nutrient concentrations (N, P = 1/10 N)(mean ± SD) (A) Changes in DOC/NPP ratio (%) under two-gradient treatments. (B) Changes in hourly dissolved oxygen release rate (mg O_2_·L^−1^·h^−1^) under two-gradient treatments. (C) Changes in nitrogen concentration under two-gradient treatments (mixed-effects model, REML). (D) Changes in phosphorus concentration under two-gradient treatments (mixed-effects model, REML). Bars represent mean values, the error bars indicate standard deviation (mean ± SD). Two-way ANOVA was used to test the effects of light, nutrient and their interaction in panels A-B, with post hoc tests corrected by Sidak (A) and Tukey (B) correction. REML analysis was used to test fixed effects (row factor, column factor and their interaction) in panels C-D, followed by Tukey’s multiple comparisons. Different lowercase letters indicate significant differences in Tukey’s post hoc multiple comparisons (*P* < 0.05), while the same letter indicates no significant difference.

The dissolved oxygen release rate (Fig.3b) exhibited a significant interaction. Two-way ANOVA analysis showed that the light×nutrient interaction was significant (*F*(1, 12) = 12.26, *P* = 0.0044), the main effect of nutrients was also significant (*F*(1, 12) = 30.02, *P* = 0.0001), while the main effect of light was not significant (*F*(1, 12) = 0.0350, *P* = 0.8547). From the marginal mean, the overall mean values of standard light and high light were close (0.1845 vs 0.1867), but the overall mean value of Std nutrients was higher than that of High nutrients (0.2176 vs 0.1535). Tukey’s post-hoc comparison further showed that under standard nutrient conditions, increasing light intensity (Std light vs High light) showed a downward trend but did not reach significance (*P* = 0.0924); while under high light conditions, high nutrients were significantly lower than standard nutrients (Std nutrients-High light vs High nutrients-High light: mean difference = −0.1050, *P* = 0.0002). In addition, standard nutrients – standard light was significantly higher than high nutrients-high light (*P* = 0.0130), and standard nutrients-high light was also significantly higher than high nutrients-standard light (*P* = 0.0082), indicating that the effect of nutrient levels on oxygen production response was dependent on light background.

Regarding the nutrient-related indicators, for ammonium concentration (Fig.3c), the restricted maximum likelihood (REML) mixed-effects model analysis showed that the row factor, column factor and their interaction were all significant (row factor: *F*(1, 23) = 70.47, *P* < 0.0001; column factor: *F*(1.113, 8.531) = 93.38, *P* < 0.0001; interaction: *F*(1.432, 10.98) = 22.05, *P* = 0.0003).

Tukey’s post-hoc comparison showed that the initial ammonium concentration was significantly higher under high nutrient conditions than under standard nutrients (e.g., Initial: Std light-Std nutrients vs Initial: Std light–High nutrients, *P* = 0.0226; Initial: Std light-Std nutrients vs Initial: High light-High nutrients, *P* = 0.0078), and a significant difference still existed between standard nutrients and high nutrients at the Final stage (e.g., Final: Std light-Std nutrients vs Final: Std light-High nutrients, *P* = 0.0100). Overall, the model-predicted mean value of Initial was higher than that of Final (4.190 vs 1.452), with a difference of 2.738 (95% CI: [2.063, 3.413])

For total phosphorus concentration (Fig.3d), significant main effects and interaction were also observed (REML mixed-effects model: row factor significant, *F*(1, 23) = 19.91, *P* = 0.0002; column factor especially significant, *F*(1.084, 8.308) = 51.87, *P* < 0.0001; and row×column interaction significant, *F*(1.069, 8.912) = 9.912, *P* = 0.0124). Tukey’s multiple comparison showed that the initial total phosphorus concentration was significantly higher in the high light-high nutrient combination than in the standard light-standard nutrients (*P* = 0.0150), and at the final stage, phosphate was significantly higher in high nutrients than in standard nutrients (e.g., Final: Std light-Std nutrients vs Final: Std light-High nutrients: *P* = 0.0045; Final: High light-Std nutrients vs Final: Std light-High nutrients: *P* = 0.0056). The model-predicted mean value also showed that Initial was higher than Final (0.2385 vs 0.1256), with a difference of 0.1129 (95% CI: [0.06056, 0.1652]).

### 3.3 Short-term day-night alternating culture experiment

There was a significant linear relationship between the dry weight and wet weight of *Z. marina* (Fig.4a). The linear regression equation was fitted as: Y = 0.1394X - 0.03088 (Y = dry weight, X = wet weight), with a high goodness of fit (*R*^2^ = 0.9800, S_y.x_ = 0.03116, n = 68). The regression slope was significantly different from 0 (*F*(1,66) = 3233, *P* < 0.0001), indicating that we can effectively predict dry weight using wet weight. The 95% CI for the slope was [0.1345, 0.1443], and the 95% CI for the intercept was [-0.04430, −0.01747].

**Fig. 4.**
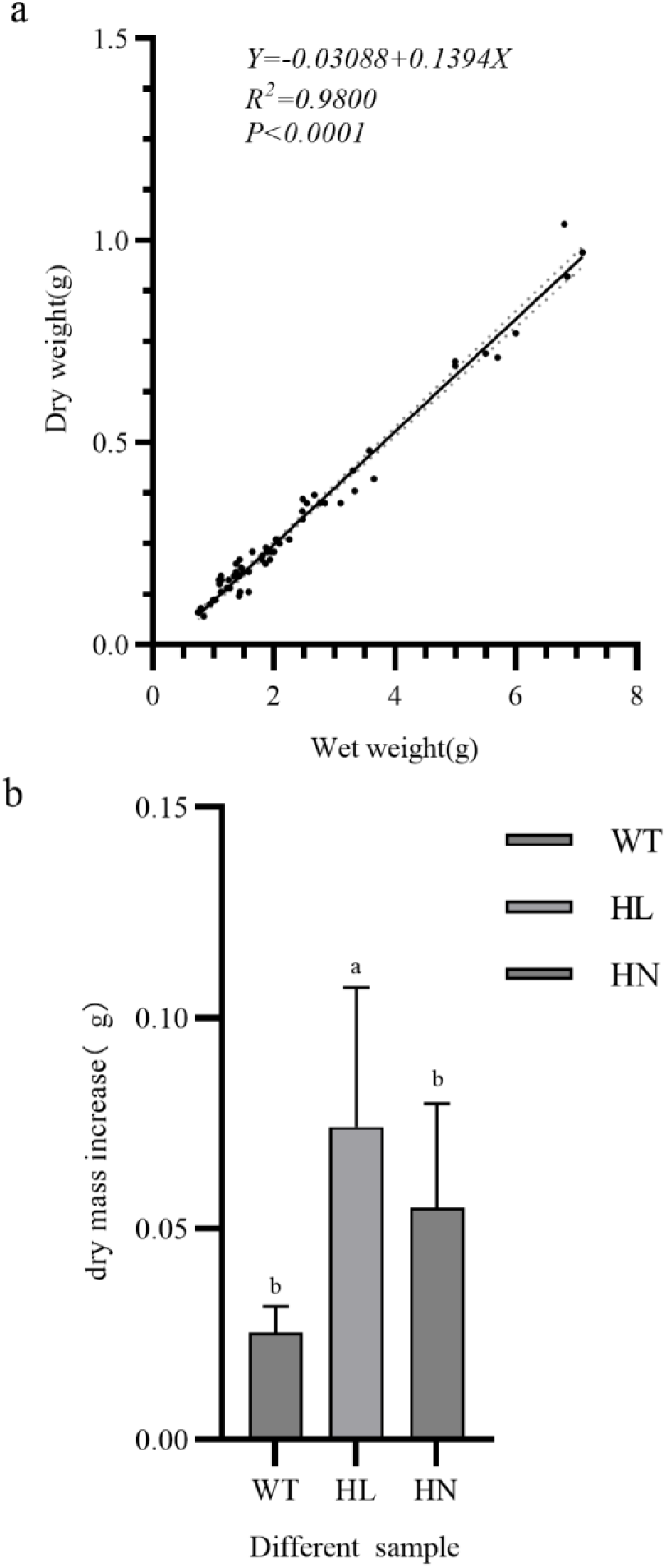
Biomass indicators of *Z. marina*: conversion between dry and wet weight and changes in dry weight before and after cultivation. (A) Linear regression relationship between dry weight and wet weight of *Z. marina* (n=68). The fitted line and its 95% CI are shown. (B) Differences in dry weight of Z. marina before and after cultivation under different treatments (dry weight increase; mean ± SD). One-way ANOVA was used for comparison among groups, followed by Dunnett’s multiple comparisons (with WT as control) to test differences between each treatment and the control. Different lowercase letters indicate significant differences in Tukey’s post hoc multiple comparisons (*P* < 0.05), while the same letter indicates no significant difference.

There were significant differences in the dry weight change of *Z. marina* before and after cultivation among different treatments (Fig.4b; one-way ANOVA: *F*(2,12) = 5.202, *P* = 0.0236, *R*^2^ = 0.4644). The Brown-Forsythe test for homogeneity of variances showed no significance (*P* = 0.1278), while the Bartlett test indicated that there was a difference in variances (*P* = 0.0256). Therefore, pairwise comparisons were primarily used for result interpretation. Dunnett’s multiple comparisons with WT as the control showed that the dry weight increase in the HL treatment was significantly higher than that in WT (WT vs HL: mean difference = −0.04879, 95% CI: [-0.08692,-0.01066], adjusted *P* = 0.0140; WT mean = 0.02537, HL mean = 0.07416; n = 5 per group). In contrast, the difference between HN and WT was not significant (WT vs HN: mean difference = −0.02955, 95% CI: [-0.06769, 0.008581], adjusted *P* = 0.1328; HN mean = 0.05492; n = 5 per group).

Under the three treatments: WT, HL, and HN, the variables in Fig.5a exhibited a nonlinear relationship with time, and a quadratic polynomial was used for fitting. The fitting results showed that WT had a high goodness of fit (*R*^2^ = 0.6995), HN also achieved a relatively good fit (*R*^2^ = 0.5790), while HL had a low goodness of fit (*R*^2^ = 0.1972), indicating that the time variation patterns of the variables may differ among different treatments.

**Fig. 5.**
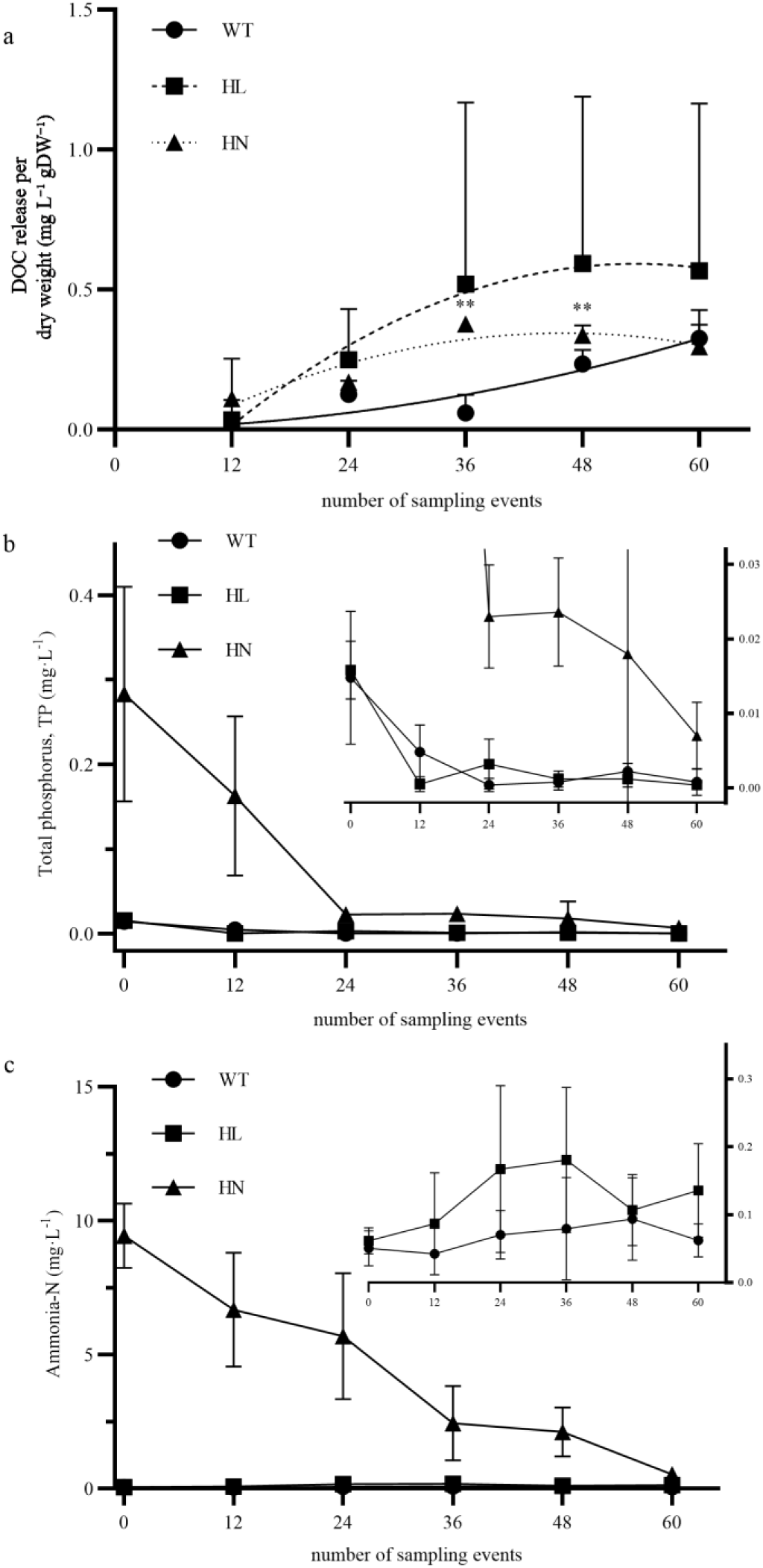
Temporal dynamics and treatment differences in fluxes and nutrient (P, N) under short-term diurnal alternating culture experiment (mean ± SD) (A) Temporal trajectory of variables under WT, HL, and HN treatment, fitted with a quadratic polynomial model (y = y_0_ + ax + bx^2^). (B) Day-night temporal variation in phosphorus concentration (P, expressed as PO_4_ ^3-^) under different treatments. (C) Day-night temporal variation in nitrogen concentration (N, expressed as NH_4_^+^) under different treatments. Data in panels B and C were analyzed using REML analysis, to test the effects of time, treatment and their interactions, followed by simple effect analyses among time points within each treatment. Multiple comparisons in panel A were corrected with Sidak correction, while significance is indicated by *P* < 0.05 and \**P* < 0.01 (comparisons between HN and WT). For nutrient analyses, multiple comparisons were corrected with Sidak (B) and Tukey (C) corrections. Different lowercase letters indicate significant differences in Tukey’s post hoc multiple comparisons (*P* < 0.05), while the same letter indicates no significant difference.

A REML mixed-effects model analysis of the DOC diel flux data showed that time (row factor) had a significant effect on flux (*F*(1.130, 4.521) = 10.19, *P* = 0.0266), whereas treatment (column factor: WT, HL, and HN) had no significant effect (*F*(1.006, 4.025) = 1.604, *P* = 0.2740). The interaction between time and treatment was also not significant (*F*(1.499, 5.995) = 1.667, *P* = 0.2585). Further in-row post hoc simple effects tests with Sidak correction revealed that treatment differences occurred only at specific time points. For example, WT was significantly lower than HN at the third time point (mean difference = −0.3166, *P* = 0.0010), and WT was also significantly lower than HN at the 4th row (mean difference = −0.1036, *P* = 0.0087); no significant differences were observed between WT, HL, and HN at all other time points (*P* > 0.05). Overall, the diel variation of flux was mainly driven by time-related factors, and the treatment effect was not obvious at most time points, with significant differences only observed between HN and WT at individual time points.

The REML analysis of diel phosphates’ variation data showed that the main effect of time, the main effect of treatment, and the time × treatment interaction were all extremely significant (time: *F*(1.430, 16.87) = 20.67, *P* < 0.0001; treatment: *F*(2, 12) = 26.15, *P* < 0.0001; time× treatment: *F*(10, 59) = 15.86, *P* < 0.0001), indicating that the temporal dynamics of phosphates differed significantly under different treatments. Further in-column post hoc simple effect tests (comparison of time points within each treatment, Sidak correction) showed that only a small number of time point differences were detected in the WT treatment (e.g., Row 3 vs Row 5: *P* = 0.0126), while multiple time point differences with Row1 were observed in the HL treatment (e.g., Row 1 vs Row 4: *P* = 0.0086; Row 1 vs Row 5: *P* = 0.0247; Row 1 vs Row 6: *P* = 0.0137), and no significant overall time point differences were found in the HN treatment (most comparisons *P* > 0.05).

Diel ammonia nitrogen concentration variation also showed significant temporal dependence and treatment differences. The REML analysis showed that the main effect of time, the main effect of treatment, and the time×treatment interaction were all extremely significant (row factor: *F*(2.604, 30.20) = 38.15, *P* < 0.0001; column factor: *F*(2, 12) = 75.64, *P* < 0.0001; interaction: *F*(10, 58) = 39.89, *P* < 0.0001). Post-hoc simple effect tests (Tukey’s test) showed that no significant differences were observed between most time points in the WT and HL treatments (most comparisons *P* > 0.05), while significant differences were observed between multiple time points in the HN treatment with large variation amplitudes (e.g., Row 1 was significantly higher than Row 3–Row 6: *P* = 0.0165, 0.0018, < 0.0001, 0.0002; Row 2 was also significantly higher than Row 5 and Row 6: *P* = 0.0344, 0.0142), indicating that the diel variation of nitrogen content under HN conditions was more intense and showed a staged downward trend.

### 3.4 Differential Expression and Pathway Enrichment

Based on transcriptome differential analysis, with the threshold set as |log2FC| > 1 and Padj < 0.05, the magnitude and number of differentially expressed genes between HL and WT were relatively small (Fig. 6b). Among the significantly differentially expressed genes, downregulation was dominant, with representative genes including SBT4-like, KIC, IP5P5-like, NRAMP2/DMT1 (ion transport-related) and others; upregulated genes were relatively few, and only DMR6/DLO and others were observed to be upregulated under HL conditions (Fig.8B). This result indicated that HL mainly triggered a limited range of transcriptional responses relative to WT, and was more inclined to inhibit partial transport/protein processing-related genes.

**Fig 6.**
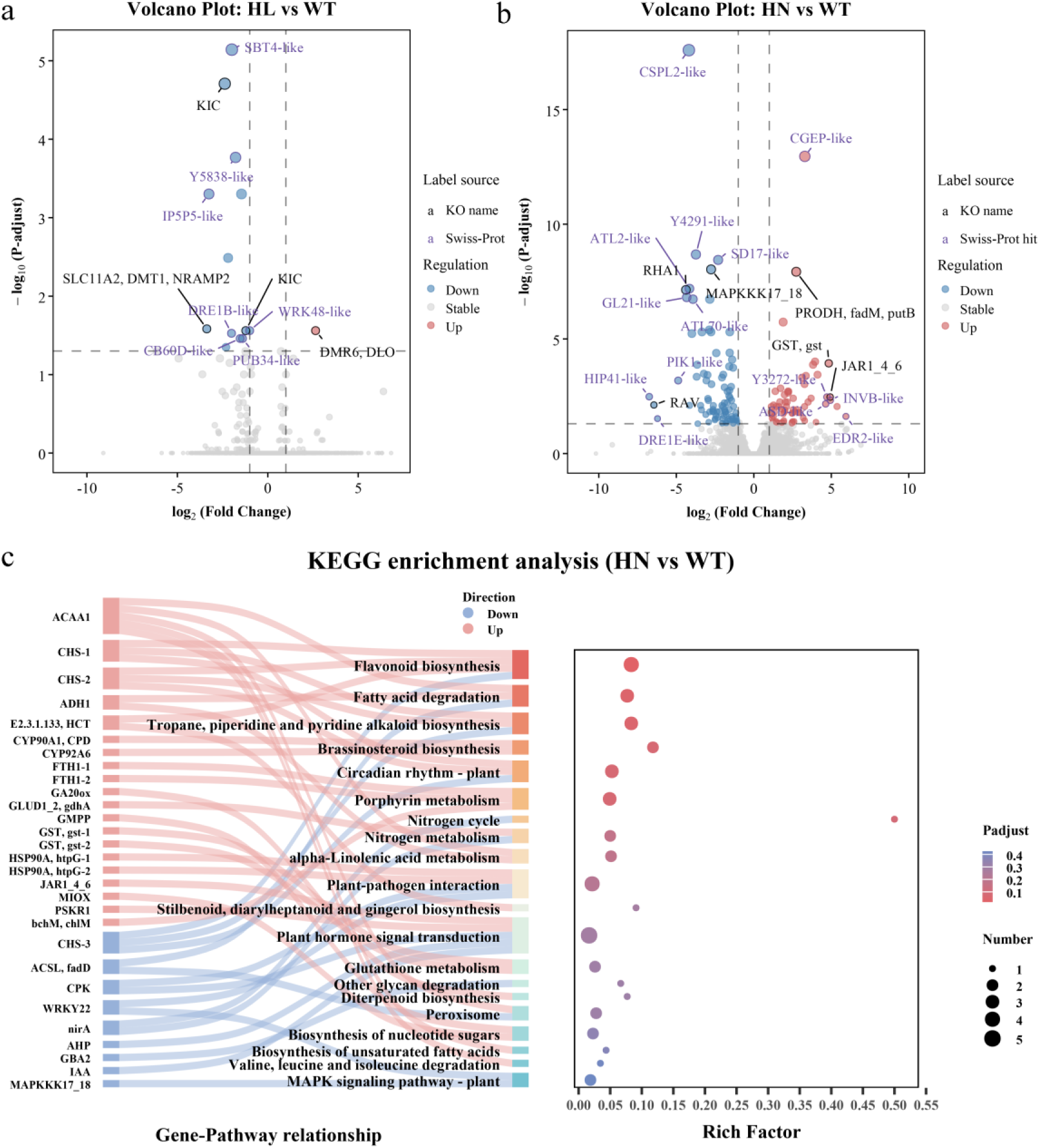
Differential expression and pathway enrichment analysis induced by high nutrient and high light treatments. (A) Volcano plot of differentially expressed genes in HL(high nutrient) vs. WT (control). The x-axis shows log_2_ (Fold Change), and the y-axis shows −log_10_ (Padj). Dashed lines represent the thresholds (|log_2_FC| = 1 and Padj = 0.05). Red, blue, and gray dots represent upregulated, downregulated, and non-differentially expressed genes, respectively. Gene labels are from KO (black) and Swiss-Prot (purple) as mentioned in the legend. (B) Volcano plot of differentially expressed genes in HN (high light) vs. WT. The definitions of axes, thresholds, and color definitions are the same as in (A). (C) KEGG enrichment results of differentially expressed genes between HN and WT. The left panel shows the gene-pathway network (line colors indicate the expression direction of genes in HN relative to WT: upregulated or downregulated). The right panel shows the enrichment bubble plot; the x-axis represents the Rich factor, bubble size represents the number of differentially expressed genes enriched in the pathway, and bubble color represents the adjusted significance level (Padj).

In contrast, a significant expression reprogramming was observed between HN and WT (Fig. 6a). The volcano plot showed that HN treatment simultaneously induced the upregulation and downregulation of multiple genes: among the upregulated genes, GST (glutathione S-transferase), PRODH/fadM/putB (proline metabolism-related genes), GA20ox (gibberellin biosynthesis-related) and others exhibited large fold changes; among the downregulated genes, CSPL2-like (casp-like), multiple receptor-like kinase/signal pathway-related genes (such as MAPKKK17_18, SD17-like, Y4291-like) and ATL family RING finger proteins (such as ATL2-like, ATL70-like) showed a significant downward trend (Fig.6). Overall, compared with WT, HN presented an expression profile characterized by the upregulation of metabolism and stress resistance-related genes, and the downregulation of partial signal transduction and receptor-like kinase-related genes.

To further analyze the functional orientation of differentially expressed genes under HN treatment, KEGG enrichment analysis was performed on the differential genes in HN vs WT (Fig.6c). The results showed that the differentially expressed genes were significantly enriched in metabolic pathways such as Flavonoid biosynthesis (Padj≈0.018), Fatty acid degradation (Padj≈0.042) and Tropane/piperidine/pyridine alkaloid biosynthesis (Padj ≈ 0.050); meanwhile, pathways including brassinolide biosynthesis, Plant circadian rhythm, Porphyrin and metabolism, Nitrogen cycle/nitrogen metabolism also showed an enrichment trend (Fig.6c). In addition, pathways related to signal regulation and stress response (such as Plant hormone signal transduction, MAPK signaling pathway), as well as antioxidant metabolic pathways such as Glutathione metabolism and Peroxisome were also detected. This suggested that under high nutrient treatment, Z. marina may drive the observed physiological and phenotypic changes by regulating secondary metabolism and lipid metabolism, and coordinating hormone and stress signaling pathways (Fig.6c).

## 4. DISCUSSION

### 4.1 Both light and nutrients affect the partitioning of fixed carbon to DOC in Z. marina, but with distinct mechanisms

In this study, we systematically compared the effects of light intensity and nutrient concentration on DOC release and its partitioning characteristics in *Z. marina* using single-factor gradient experiments, two-factor orthogonal experiments, and Short-term day-night alternating culture experiments under controlled laboratory conditions. The underlying mechanisms were further explored by TEM observation and transcriptome analysis. Overall, both light and nutrients affected the partitioning ratio of fixed carbon released as DOC (DOC/NPP) in *Z. marina*, but their regulatory mechanisms differed.

Notably, according to our results, an increase in DOC/NPP can arise from either enhanced DOC release or reduced NPP. Therefore, this index mainly reflects changes in the relative partitioning of fixed carbon released into the dissolved phase, and cannot be simply used to determine the specific mechanism of DOC production. Based on the present results, we propose that this phenomenon can be interpreted within a “source–sink regulation” framework: light mainly regulates the supply of carbon assimilation, whereas nutrients more strongly influence the allocation of fixed carbon among growth, metabolic maintenance, and release in dissolved form, with the latter exerting a stronger effect(Paul and Foyer, 2001; Rosado-Souza et al., 2023).

Specifically, high light may enhance “source” supply, leading to excess fixed carbon being exuded as DOC when growth demands are basically satisfied. In contrast, high nutrients may suppress “sink” utilization and induce stress-related metabolic reprogramming, diverting more fixed carbon toward antioxidant, defense, and secondary metabolic processes, ultimately increasing the proportion of DOC in NPP. Nevertheless, this interpretation bases on phenomenological and indirect evidence, and requires further experimental validation.

### 4.2 High light may increase the proportion of DOC efflux by enhancing carbon assimilation and inducing cellular stress

Light gradient experiment showed that DOC/NPP in *Z. marina* increased significantly while PPFD = 350, consistent with previous studies(Kaldy, 2012), which also supports the view that high light promotes carbon fixation and makes excess fixed carbon release in dissolved form(Fogg, 1983). Notably, the oxygen evolution rate did not peak when PPFD = 350, but at PPFD = 200, suggesting that *Z. marina* may have experienced photoinhibition or altered energy use efficiency under ultra-high light conditions(Khan et al., 2025; Song et al., 2021). Under such conditions, when assimilated carbon exceeds the demand for growth and metabolism, part of the newly fixed carbon may be diverted to DOC efflux, resulting in elevated DOC/NPP(Adams et al., 2016). However, the number of differentially expressed genes (DEGs) under high light was relatively low, and no significantly enriched pathways were detected, implying that the short-term high-light response in *Z. marina* may involve rapid changes in both the cellular structural and physiological levels rather than substantial transcriptomic reprogramming at the sampling time point, which suggests that short-term defense may rely more on existing physiological regulation and protein repair rather than large-scale transcriptional responses(Dziubek et al., 2024; L. Li et al., 2022). Therefore, we used transmission electron microscopy to examine the leaf structure of Eelgrass at the end of a Short-term day-night alternating culture experiment and found significant structural changes compared to the wild-type(Fig.7): the high-light treatment exhibited swollen chloroplasts, reduced or blurred grana structures, ruptured chloroplast outer membranes, and mitochondrial vacuolization, indicating that membrane systems and energy-related organelles were under stress, which may increase the likelihood of DOC efflux or leakage(Song et al., 2021; Tang and Zhu, 2023). Furthermore, passive DOC efflux caused by structural damage may be quicker than active transcriptional responses, which could explain why DOC/NPP increased under high light without significant DEGs detected(Kaldy, 2012; Yoshioka-Nishimura, 2016). It must be emphasized that both TEM and transcriptomic results are indirect evidence and cannot directly demonstrate the specific mechanism of DOC efflux.

**Fig. 7.**
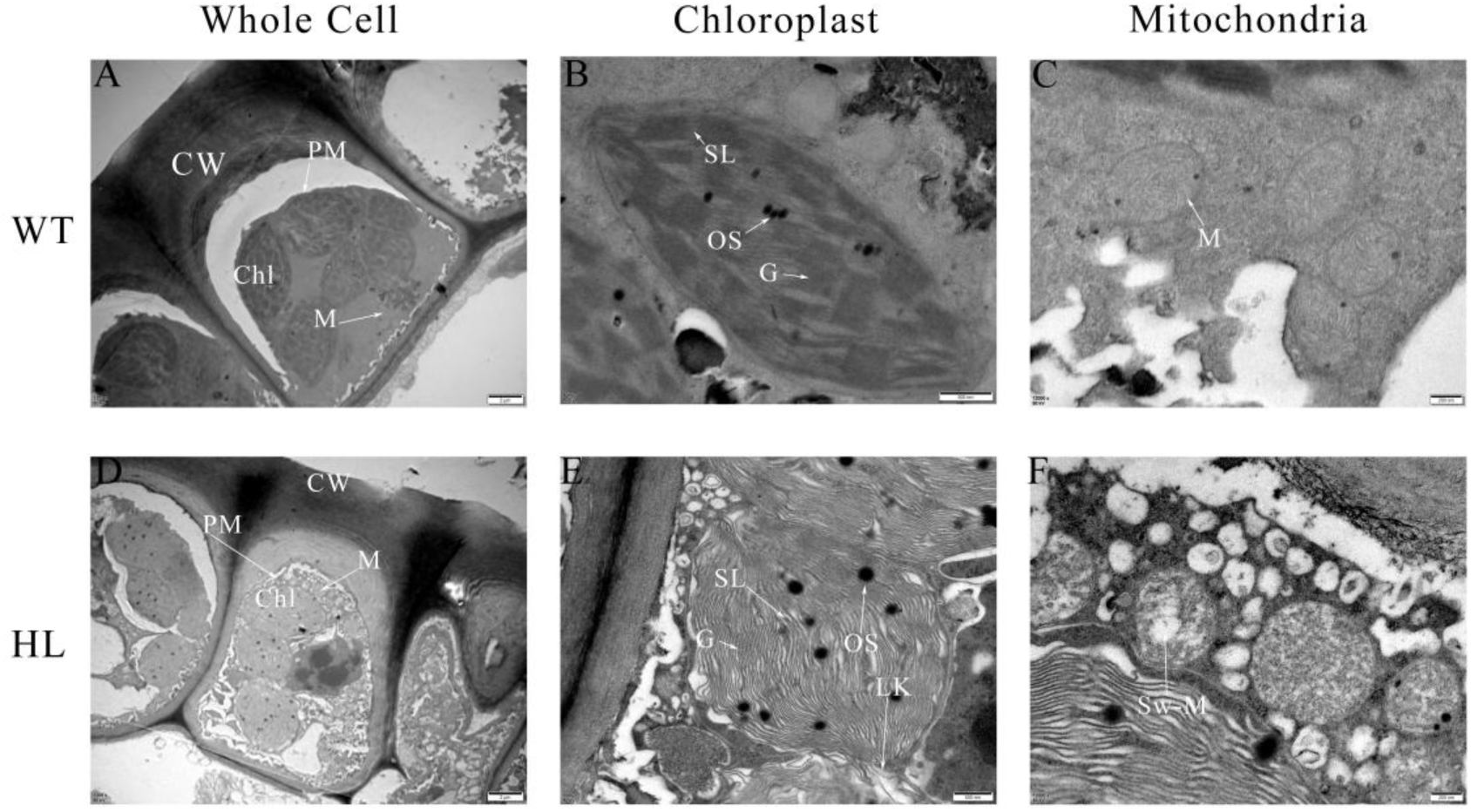
TEM micrographs showing leaf epidermal cell morphology, chloroplast ultrastructure, and mitochondrial ultrastructure of *Z. marina* under three treatments. (a–c) WT (wild type): epidermal cell (a), chloroplast (b), and mitochondrion (c). (d–f) HL (high light): epidermal cell (a), chloroplast (b), and mitochondrion (c). Scale bars: 2 μm (a, d), 500 nm (b, e), 200 nm (c, f). At least 10 fields were observed per plant, and images are representative of n = 5 biologically independent plants per treatment. Images were photographed under a transmission electron microscope (JEOL, JEM-1400F, Japan) at an accelerating voltage of 80 kV. Ultra-thin slices (∼70 nm) were prepared as mentioned in Section 2.4.3. Brightness and contrast were uniformly adjusted across all images, and no structural features were added or removed. Abbreviations: CW, cell wall; PM, plasma membrane; Chl, chloroplast; M, mitochondrion; SL, stromatolites lamella; OS, osmiophilic globule; G, grandma; Sw-M, swollen mitochondrion; LK, matrix leakage

TEM observations provided indirect evidence: Based on the above ultrastructure, physiological, and transcriptomic results, we further constructed a conceptual model of DOC efflux under high light (Fig.8). This model indicates that the elevation of DOC/NPP by high light may not only result from enhanced carbon fixation, but also from increased passive efflux associated with altered membrane permeability and metabolic imbalance following organelle damage.

**Fig. 8.**
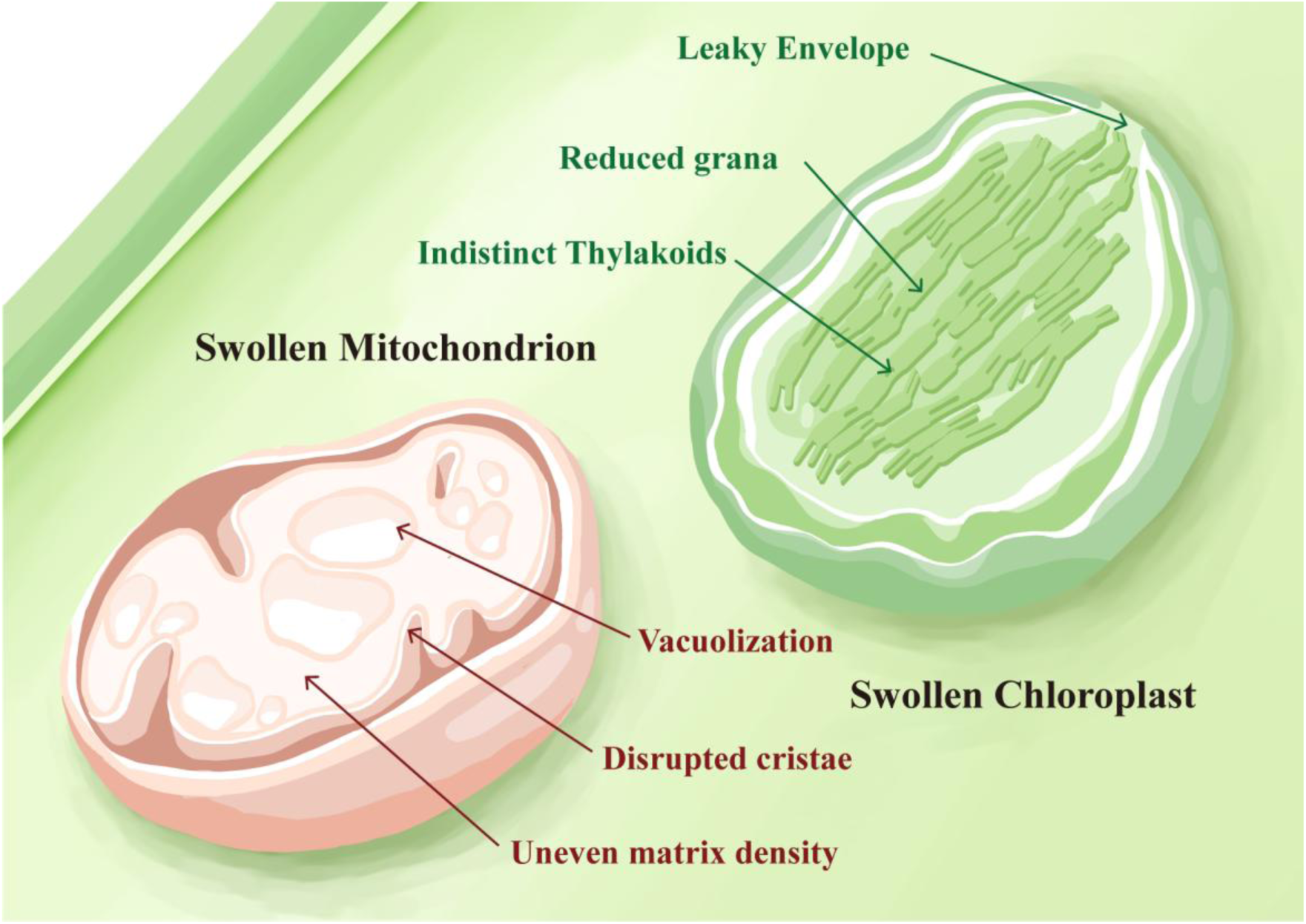
Diagram of the response pattern of epidermal cell structure in *Z. marina* leaves under high-light treatment. This model is constructed based on the observations of the ultrastructure of epidermal cells of *Z. marina* leaves after HL treatment in this study, aiming to summarize the possible processes of organic carbon allocation changes in *Z. marina* under the treatment. Ultrastructural observations of cells reveal that significant alterations occur in chloroplasts and mitochondria of *Z. marina* cells under high light conditions. Some mitochondria swell an undergo vacuolization; in the chloroplasts, the boundaries between grana and stroma thylakoids become blurred, meanwhile some partial leakages appear at the chloroplast envelope.

### 4.3 Nutrients dominate the variation in the proportion of DOC efflux and may act via stress-driven metabolic redistribution

Compared with light, nutrients exerted a stronger effect on DOC/NPP. In the single-gradient experiment, DOC/NPP increased from10.38% to 37.99%-52.41% and tended to plateau. In the two-factor orthogonal experiments, high nutrients elevated its mean value from 40.33% to 92.83%. These results generally support the diffusion mechanism hypothesis that nutrients dominate DOC efflux(Bjørrisen, 1988). However, our results differed from those of previous nitrate enrichment experiments on *Thalassia hemprichii*(Z. Jiang et al., 2022), which may be related to differences in nitrogen form and associated metabolic costs(Jiménez-Ramos et al., 2022b). The ammonium (NH_4_ ^+^) used in this study can be rapidly assimilated, but excessive uptake often induces physiological stress in seagrass(F. Jiang et al., 2017). Consistent with this, oxygen production rate generally decreased with increasing nutrient concentration. In the two-factor orthogonal experiments, nutrients showed a significant main effect on oxygen production and a significant interaction with light.

Therefore, the phenomenon of increased DOC/NPP accompanied by decreased oxygen production more likely reflects a metabolic redistribution caused by high nutrients, rather than a simple increase in DOC efflux caused by enhanced photosynthesis(Touchette and Burkholder, 2000). In this study, KEGG enrichment analysis showed that most enriched pathways contained a small number of matched genes, and the overall Rich Factor was relatively low. This pattern is consistent with the trends observed in previous studies on transcriptome, which also focused on the short-term responses of adult *Z. marina* leaves to environmental factors(Yan et al., 2023, 2024). This feature may first reflect that under short-term, non-extreme environmental stress, differentially expressed genes exhibit dispersed responses throughout the metabolic network, rather than being concentrated in a few core pathways. In addition, *Z. marina* has undergone substantial genome remodeling during its evolutionary adaptation to the marine environment(Olsen et al., 2016). For such non-model organisms, current KEGG annotation and pathway classification mainly rely on existing reference gene sets and homologous function inference. Consequently, some *Z. marina*-specific genes or functional genes that differ greatly from known model plant genes may not be accurately mapped to established ways. Transcriptomic results further revealed that HN treatment triggered more pronounced molecular responses, involving pathways such as antioxidant defense, nitrogen metabolism, hormone signaling, lipid and energy metabolism, and flavonoid metabolism, which suggests that under high nutrient conditions, *Z. marina* may increase the proportion of DOC allocated to fixed carbon through stress responses and carbon flow redistribution(Brunn et al., 2025; Zou and Xu, 2025). Overall, nutrient changing regulates the fate of fixed carbon in *Z. marina* more strongly than light, and more directly affected the partitioning of fixed carbon among biomass accumulation, metabolic maintenance, and DOC efflux. According to transcriptomic consequence, a hypothetical mechanistic model of metabolic pathways under HN conditions was constructed(Fig.9)。This model indicates that the elevation of DOC/NPP under high nutrient treatment may be mainly associated with increased nitrogen assimilation demand, enhanced oxidative stress, and the consequent metabolic reprogramming.

**Fig. 9.**
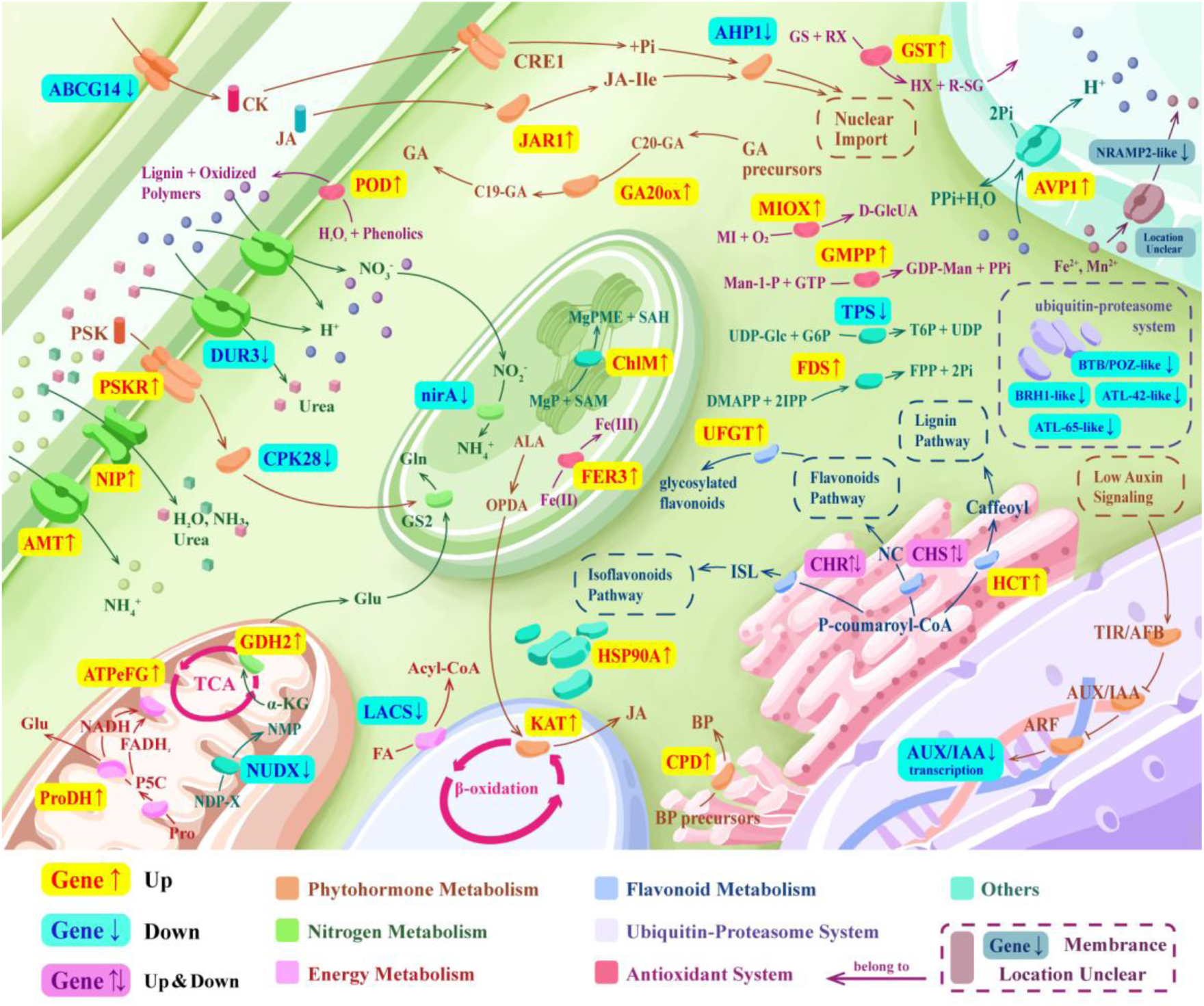
Diagram of cellular metabolic response pattern in *Z. marina* leaf cells under high-nutrient treatment. This model is constructed based on the transcriptomic analysis and ultrastructural observation results of *Z. marina* leaf cells after HN treatment in this study, aiming to summarize the possible synergetic regulatory processes within *Z. marina* cells under the treatment. Transcriptomic analysis revealed that the effects of HN treatment could be broadly categorized into seven major response groups. These included enhanced nitrogen uptake and assimilation (grass green; e.g., upregulation of AMT, NIP, GDH2, and downregulation of DUR3 and nirA), accompanied by strengthened antioxidant defense and metal homeostasis (rose pink; e.g., upregulation of POD, FER3, GST, MIOX, GMPP, and NRAMP). In addition, plant hormone signaling was extensively remodeled (apricot orange; e.g., upregulation of JAR1, GA20ox, CPD, and PSKR, and downregulation of ABCG14, AHP1, AUX/IAA, and CPK28), along with adjustments in energy and lipid metabolism (light magenta; e.g., upregulation of ProDH and ATPeFG, and downregulation of LACS). Changes were also observed in flavonoid secondary metabolism and cell wall-related processes (light blue; e.g., upregulation of HCT and UFGT, and differential expression of CHR and CHS).Meanwhile, ubiquitination-related genes (lavender white; including BTB/POZ-like, BRH1-like, ATL-42-like, and ATL-65-like) were generally downregulated, suggesting a potential reduction in protein degradation under stress conditions. Additionally, several unclassified genes (mint green; e.g., upregulation of AVP, HSP90A, ChlM, and FDS, and downregulation of TPS and NUDX) were also involved. Collectively, these results suggest that high nutrient conditions may induce oxidative stress and trigger extensive hormonal and metabolic reprogramming in *Z. marina*, redirecting fixed carbon from growth and biomass accumulation toward defense, secondary metabolism, and dissolved organic carbon (DOC) release, thereby increasing the proportion of DOC allocation within net primary production (NPP).

Notably, these models are inferred primarily from transcriptomic results and still require further verification using key enzyme activities, targeted metabolomics, and isotope tracing techniques.

### 4.4 Energy metabolic responses are asynchronous with changes in DOC partitioning

Two-factor orthogonal experiment showed no significant interaction effect on DOC/NPP, suggesting that under our experimental conditions, the proportion of DOC efflux is more likely an additive response dominated by nutrients and secondarily modulated by light. In contrast, oxygen production exhibited a significant light × nutrient interaction, indicating that energy metabolic processes are more sensitive to nutrient conditions(Jiménez-Ramos, Villazán, et al., 2022). These results demonstrated that DOC efflux was not necessarily synchronized with photosynthetic intensity, but mainly reflected the partitioning of fixed carbon among different fates. By comparison, oxygen production more directly represents the net balance between photosynthetic production and respiratory consumption(Paul and Foyer, 2001; Barrón et al., 2014; Hansen et al., 2022).

### 4.5 Short-term day-night alternating culture experiment further supports differential regulation of fixed carbon allocation pathways by light and nutrients

Short-term day-night alternating culture experiment further supported the differential regulation of fixed carbon allocation pathways in *Z. marina* by different environmental factors. Dry weight results showed that the dry weight increment under HL was significantly higher than that under WT, while no significant difference was observed between HN and WT, which suggests that under high lights part of the newly fixed carbon was allocated to biomass accumulation in addition to DOC efflux(Alcoverro et al., 1999; Zimmerman et al., 1997), while this inference still requires further experimental verification. Combined with the elevated DOC/NPP under high light, we proposed that increased light mainly enhances carbon assimilation at the supply side, directing fixed carbon toward both growth and DOC release. Combined with the elevated DOC/NPP under high light, we proposed that increased light mainly enhances carbon assimilation at the supply side, directing fixed carbon toward both growth and DOC release. In contrast, high nutrients increased the proportion of DOC efflux but did not significantly promote dry weighted accumulation, indicating that the associated carbon flow shift was more likely a stress-driven redistribution rather than growth stimulation.

Previous studies have revealed that seagrass responses to ammonium enrichment are concentration-dependent. Excessively high ammonium concentrations can be toxic or stressful to *Z. marina*(F. Jiang et al., 2017), inducing nitrogen metabolism and detoxification processes that do not necessarily lead to increased biomass(Liu et al., 2025; Van Katwijk et al., 1997). Meanwhile, DOC flux results indicated that its variation was mainly driven by time, with treatment effects being insignificant at most time points, except for occasional differences between HN and WT at certain stages. In terms of release patterns, the mean DOC flux in the HN group was higher than in WT during the initial take buy gradually converged at 60 h, which suggested that the stimulatory effect of nutrient enrichment on DOC efflux may be prominent mainly in the short-term response phase and gradually weakens as available nutrients in the culture system decreased. By comparison, although the HL group showed relatively high DOC flux at some time points, large within-group variability weakened the significance of the overall treatment effect.

### 4.6 Environmental factors may alter the retention and export balance of blue carbon in seagrass beds by affecting DOC allocation

During the blue carbon fixation process in seagrass beds, changes in the release proportion of DOC may affect the retention of fixed carbon in local systems and its export to surrounding water column(Gomis et al., 2025; Kindeberg et al., 2019), as DOC is a mobile carbon form. Existing studies have shown that organic carbon released from seagrass beds is not completely retained in situ, but can be exported outward through multiple pathways, among which DOC is a crucial component(Krumhansl et al., 2025). This DOC can either be rapidly utilized by microorganisms or transported with water currents to participate in larger-scale carbon cycling processes (Zhang et al., 2025).

The results of this study demonstrate that environmental factors not only affect the apparent productivity of *Z. marina*, but also alter the balance between organic carbon retention and carbon export in seagrass beds by changing the allocation proportion of DOC in fixed carbon. Therefore, it is necessary to incorporate DOC efflux and its environmental regulation into the evaluation of the blue carbon fixation capacity of seagrass beds. Future studies should further distinguish the relative contributions of leaves and roots-rhizomes to DOC release(Kaldy, 2012; Sun et al., 2025), and combine different life history stages and complex environmental scenarios closer to natural conditions to deeply evaluate the role of DOC flux in the blue carbon function of seagrass beds.

## 5. CONCLUSION

This study indicates that both light intensity and nutrient concentration significantly influence DOC release in *Z. marina*. The high-light treatment increased the DOC/NPP ratio and resulted in a rise in eelgrass biomass, accompanied by increased dissolved oxygen release. This suggests that light enhances the net primary productivity of eelgrass and facilitates the release of excess carbon from the plant. Additionally, light-induced cellular structural damage further increases DOC release.

Compared to high light, high nutrient concentrations exerted a stronger influence on the DOC/NPP ratio. Nutrient enrichment significantly increased the DOC release rate while reducing net primary productivity, and elicited a stronger transcriptomic response during short-term cultivation. In contrast, fewer genes were differentially expressed under high-light treatment. Overall, this study demonstrates that both high light and high nutrient concentrations promote the release of carbon sequestered by eelgrass in the form of DOC, but their mechanisms of action differ: high light primarily acts by enhancing carbon supply and inducing cellular structural stress, whereas high nutrient concentrations may promote DOC release through metabolic reallocation and physiological stress. These findings provide insights into the mechanisms of DOC release from seagrass beds and offer a reference for blue carbon assessment.

## 6. DATA AVAILABILITY STATEMENT

Data will be made available on request.

## 7. ACKNOWLEDGMENT

This work was supported by National Undergraduate Training Program for Innovation and Entrepreneurship (202510423024) and Fund of Observation and Research Station of Laizhou Bay Marine Ecosystem, MNR, and Shandong Key Laboratory of Marine Ecological Restoration (SAL202413)-

