## Supplementary material for "DOC Release and Physiological Response of *Zostera marina* Under Light and Nutrient Gradients": Supplyment material

**Supplement Table 1**

|  | PPFD (μmol·m-2·s-1) | N concentration (mg·L-1) | P concentration (mg·L-1) |
| --- | --- | --- | --- |
| WT | 95.12 ± 4.17 | 0.232 ± 0.087 | 0.009 ± 0.006 |
| 2.4 | 95.25 ± 12.14 | 2.623 ± 0.190 | 0.292 ± 0.018 |
| 4.8 | 95.00 ± 5.70 | 5.551 ± 0.182 | 0.567 ± 0.081 |
| 9.6 | 94.88 ± 3.97 | 8.834 ± 1.060 | 0.968 ± 0.106 |

**Supplement Table2**

|  | PPFD (μmol·m^-2^·s^-1^) |
| --- | --- |
| 50 | 49.9 ± 2.7 |
| WT | 105. ± 6.3 |
| 200 | 204.5 ± 5.1 |
| 350 | 360.5 ± 5.4 |

**Supplement Table 3**

|  | PPFD (μmol·m^-2^·s^-1^) | N concentration (mg·L^-1^) | P concentration (mg·L^-1^) |
| --- | --- | --- | --- |
| WT | 92.3 ± 14.6 | 0.111 ± 0.073 | 0.01225 ± 0.00369 |
| HL | 366.6 ± 32.1 | 0.160 ± 0.127 | 0.02175 ± 0.01135 |
| HN | 91.8 ± 5.0 | 8.57 ± 1.07 | 0.46025 ± 0.06671 |
| HLHN | 366.3 ± 13.1 | 8.42 ± 1.43 | 0.52575 ± 0.10684 |

**Supplement Table4**

|  | PPFD (μmol·m^-2^·s^-1^) | N concentration (mg·L^-1^) | P concentration (mg·L^-1^) | Wet weight (g) |
| --- | --- | --- | --- | --- |
| WT | 95.41 ± 12.98 | 0.050 ± 0.026 | 0.015 ± 0.009 | 43.59 ± 2.39 |
| HL | 300.46 ± 5.48 | 0.061 ± 0.020 | 0.016 ± 0.004 | 43.91 ± 1.93 |
| HN | 96.11 ± 10.49 | 9.44 ± 1.18 | 0.283 ± 0.121 | 44.60 ± 1.79 |
